# Identifying variants associated with dwarfism via long read sequencing of the *Ddw1* locus in winter rye

**DOI:** 10.1101/2023.04.17.537231

**Authors:** Rowena C. Haardt, Brigitta Schmiedchen, Andres Gordillo-Rodriguez, Jakob Eifler

## Abstract

Dwarfism or semi-dwarfism is an established characteristic in the European cereal market, for example semi-dwarf barley and wheat are commonplace. However, dwarf winter rye is still atypical commercially and fundamental research into the underlying genes influencing plant height is still in its relative infancy. In this study, association between two independent dwarf donors are established as *Ddw1* carriers and variants within the *Ddw1* locus are identified using long read sequencing. Here over 20,000 variants are identified as having putative association with the dwarf phenotype by using two *Ddw1* genotypes and four representative (non-dwarf) elite lines for exclusion. Consequently, the sequence data presented here provides rye breeders and researchers a resource to develop KASP markers to track the allelic status of *Ddw1* and to select for dwarf material.

## Introduction

Rye is an excellent source for the production of food, feed and bioenergy due to its outstanding tolerance against biotic and abiotic stress and its thrifty use of resources. Despite the considerable breeding progress in grain yield within the last decades, modern hybrid rye varieties are still susceptible for lodging causing the regular use of growth regulators for practical farming [8]. Since the green revolution, it has been proven that dwarf varieties in cereals provide superior yields with increased stability and less inputs [17]. However, such varieties have not yet been established across the major commercial markets in rye. In contrast to the great number of available genetic resources categorised as dwarfs or as displaying a degree of dwarfism, there are only a handful of short-straw cultivars commercialized at present. Therefore, dwarfism in rye remains an untapped resource. Despite this hesitancy to establish dwarf rye as a commonplace market feature, the potential of this phenotype has been acknowledged for many years within the realms of academia. Besides several QTL controlling plant height [13], dwarfing genes have been reported. Most of them having a recessive character and are either gibberellic acid sensitive or insensitive [15]. Among the gibberellic acid sensitive individuals, four dominant loci have been identified throughout the genome. The dominant dwarfing gene *Ddw1* (Hl / Humilus) originated from an EM-1 mutant in the germplasm collection preserved at the Vavilov Institute of Plant Industry in St. Petersburg and was originally described by Kobylyanski (1972) [6]. It was first mapped in 1996 on chromosome 5RL by using RFLP markers (Korzun et al. in 1996). The *Ddw2* locus found in a Bulgarian Mutant K10028 was described by Melz [12] and is located on chromosome 7R. Attempts have also been made to characterize the loci *Ddw3* on chromosome 1RL [16] and *Ddw4* on chromosome 3R [5].

In 2019, research into dwarfism progressed significantly with the identification that *ScGA2ox12* was found to co-segregate with the *Ddw1* locus on chromosome 5R in conjunction with increased expression in semi-dwarf individuals being observed [3]. *ScGA2ox12*, which is a gibberellic acid biosynthesis gene, is located with the identified 5.4 cM *Ddw1* locus on 5R. This locus was further described as orthologous to the Rht12 locus on chromosome 5A in wheat. In the study a set of markers were designed to support *Ddw1* tracking, some located within *GA2ox12* directly. It was suggested that *GA2ox12* may be the functional gene underlying the *Ddw1* locus, alternatively it may lead to transcriptional activation of *Ddw1*, although it is of note that at this stage, *ScGA2ox12* has not been confirmed as the candidate gene for *Ddw1*.

Observing the variation landscape within the *Ddw1* locus will enable KASP marker development for an easy and cost-effective method of tracking the *Ddw1* allelic status, as well as the flanking region status in order to reduce linkage drag.

## Results

### Phenotyping

Utilising two independent dwarf donors, three BC2 mapping populations were developed via two rounds of back-crossing to elite breeding material (non-dwarf). At the EC69 stage of development, all individuals were phenotyped based on their height in centimeters (Figure 1; Figure 2). The individuals were then designated as either dwarf or tall relative to the height segregation observed within each population.

**Figure 1.**
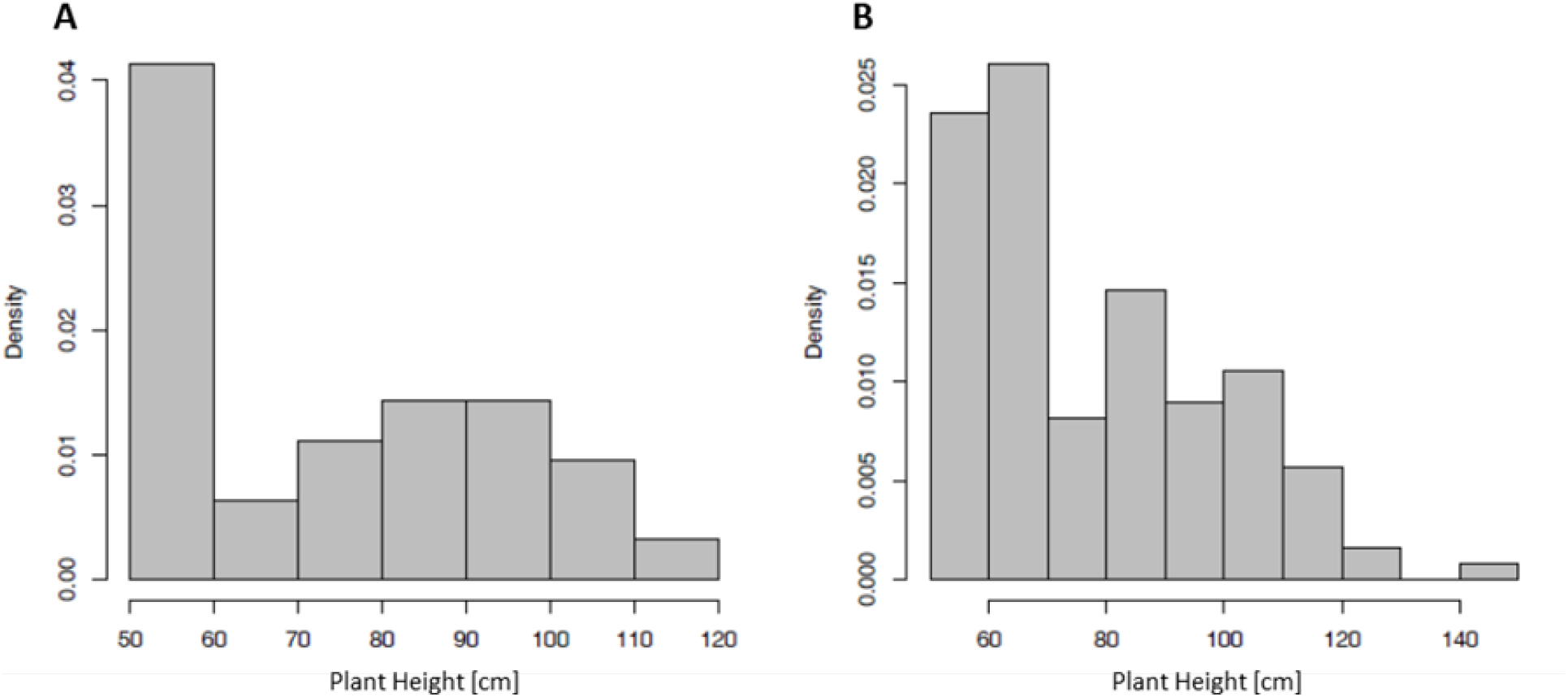
Histograms showing the segregation in plant height of two Donor A-based BC2(tall x dwarf) mapping populations. **A**: *n* = 63, *n*_*Ddw1ddw1/ddw1ddw1*_ = 29*/*34; **B**: *n* = 123, *n*_*Ddw1ddw1/ddw1ddw1*_ = 61*/*62. Here, a 1:1 dwarf to tall ratio can be observed in both populations.

**Figure 2.**
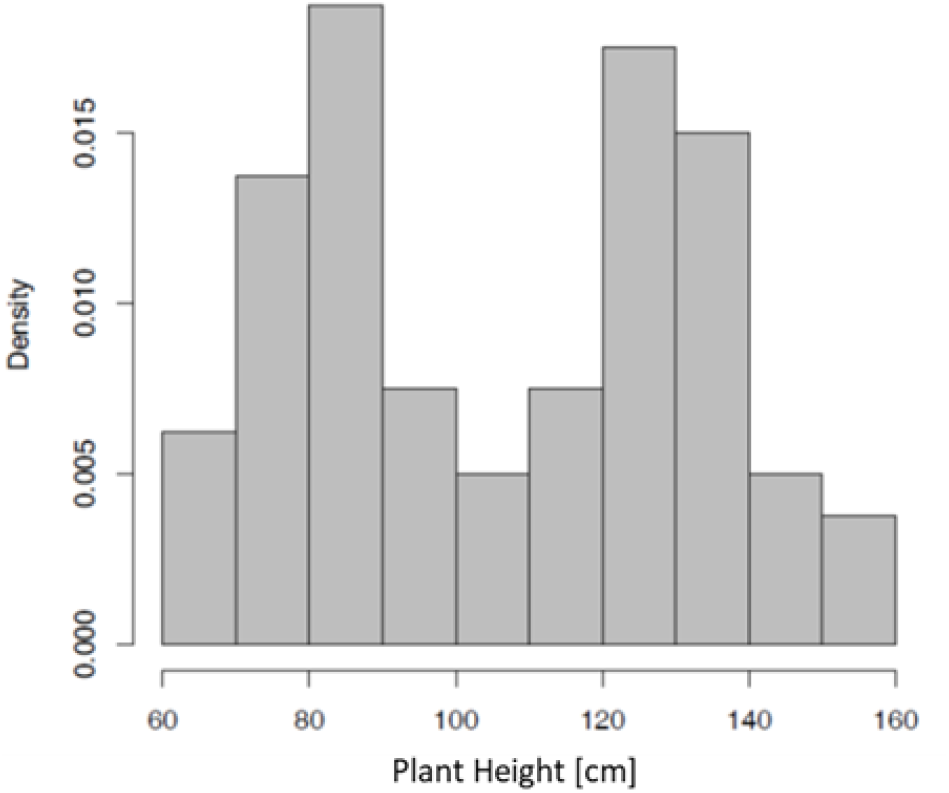
Histogram showing the segregation in plant height of Donor B-based BC2(tall x dwarf) mapping populations. *n* = 80, *n*_*Ddw1ddw1/ddw1ddw1*_ = 40*/*40. A 1:1 dwarf to tall ratio can be observed here.

Two donor A derived BC2 mapping populations, consisting of 63 and 123 individuals respectively, were selected for analysis. The ratios of *Ddw1ddw1/ddw1ddw1* observed were 1:1 in both populations (*χ*^2^, *n* = 63, *df* = 1, *p* = 0.53; *χ*^2^, *n* = 123, *df* = 1, *p* = 0.93 respectively; Figure 1). The third population with a donor B derived pedigree contained 80 individuals showed a *Ddw1ddw1* /*ddw1ddw1* ratio of 1:1 as well (*χ*^2^, *n* = 80, *df* = 1, *p* = 1; Figure 2).

### GWAS Results

Following genotyping of the donor A-derived populations using an internally designed SNP chip, a data matrix of 12,908 SNPs across the 186 individuals from the donor A-derived mapping populations was constructed. Using a significance threshold of *P* = 0.05 (*−log*_10_*P* = 5.3 with FDR correction), the GWAS analysis identified 70 significant markers associated with plant height type (i.e. dwarf = 0, uncategorized individuals between dwarf and tall measurements = 0.5, tall = 1). When using a threshold of *−log*_10_*P* = 15, which was arbitrarily designated as highly significant, GWAS analysis identified four highly significant markers associated with plant height. All significant and highly significant markers co-locate to a single hit on chromosome 5R (Figure 3).

**Figure 3.**
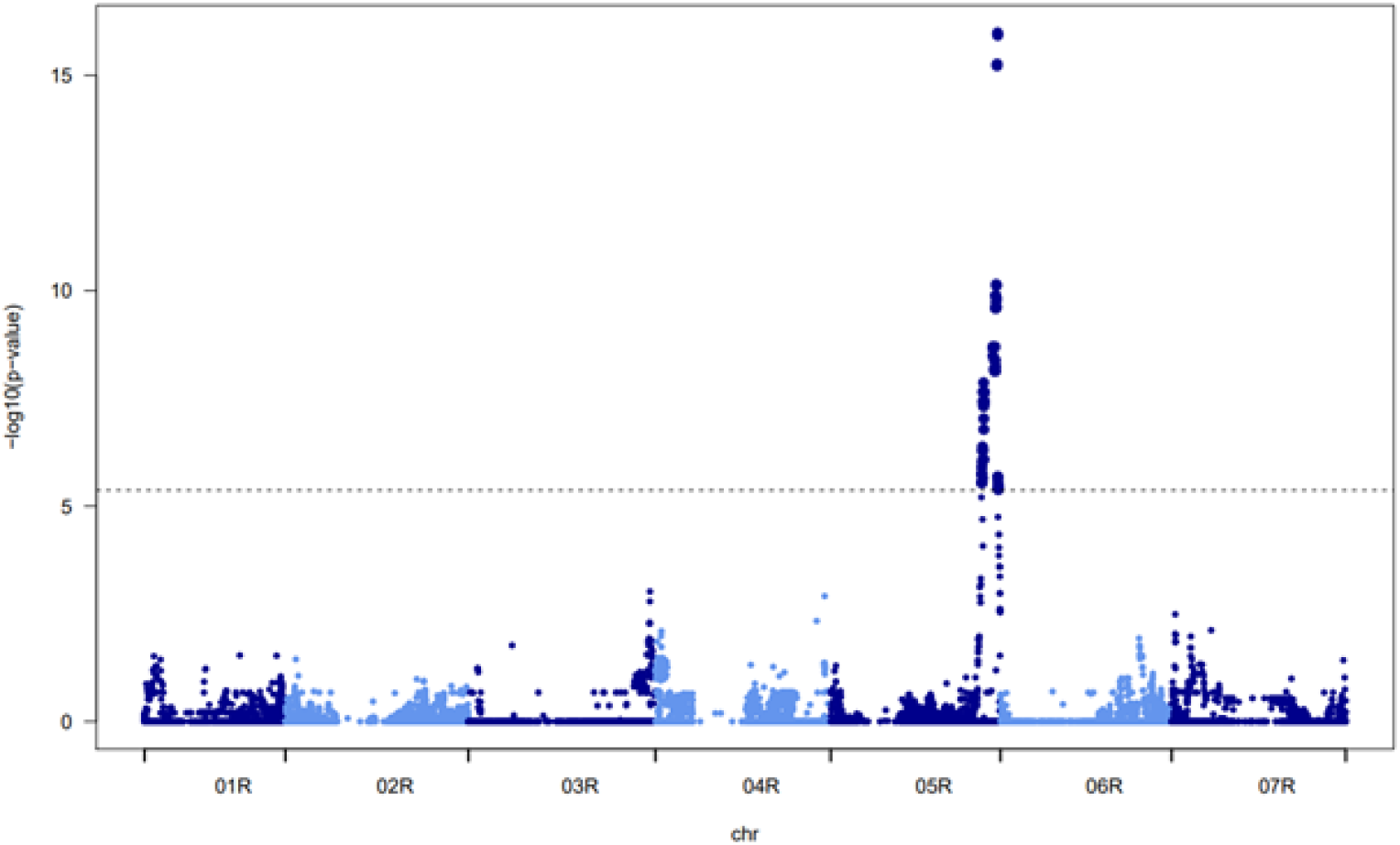
Manhattan plot of the performed GWAS. Here showing a highly significant single hit on 5R associated with plant height type (i.e. dwarf = 0, uncategorized individuals between dwarf and tall measurements = 0.5, tall = 1). Significance threshold: *−log*_10_*P* = 5.3.

The 70 significant markers determine a physical region on chromosome 5R from 779580834 bp to 865908032 bp, denoting a region of 334 SNP markers in total. The four highly significant markers, which enclose a region of 13 markers total, are located on chromosome 5R from 858846334 bp to 862353844 bp on rye Lo7 reference genome (accession number: GCA 902687465.1; Rabanus-Wallace et al. 2021; Table 1). Of these four highly significant markers, ro00281s01 was found to be within a gene model, the other three are located in intergenic sequence, according the Lo7 high confidence annotation [14]. Between the four highly significant markers, 49 high confidence gene models are present. These include four 2-oxoglutarate (2OG) and Fe(II)-dependent oxygenase superfamily proteins, two auxin canalization proteins, one cytochrome P450 family protein, an alcohol dehydrogenase, sugar transporter, putative and gibberellin 2-beta-dioxygenase.

**Table 1.** Highly significant markers identified by GWAS. This table shows marker position relative to the Lo7 reference genome, the log_10_P value, R^2^ and, if the marker SNP is located within any high confidence gene models in the Lo7 annotation, the gene model description.

| Marker | Chromosome | Position (bp) | $-\log_{10}p$-value | $R^2$ | Genic region? |
| --- | --- | --- | --- | --- | --- |
| Ro17245s01 | 5R | 858846334 | 15.22369 | 0.317 | intergenic |
| Ro01499s01 | 5R | 859366038 | 15.26131 | 0.318 | intergenic |
| Ro00281s01 | 5R | 861884664 | 15.98368 | 0.344 | genic |
| Ro00885s02 | 5R | 862353975 | 15.93049 | 0.348 | intergenic |

### Sequencing

Long read sequencing was undertaken using four elite lines and two *Ddw1* donors, representing the wild type and dwarf phenotype respectively. The two *Ddw1* donors (donor A and donor B, Table 2) were both shown to segregate for plant height when backcrossed with elite material (Figures 1 and 2). Following long read ONT sequencing 17,281,724 reads were generated with an average length of 30,332 bp. After trimming and quality control, the reads were mapped to the Lo7 reference genome (Rabanus-Wallace et al. 2021). Of these, 188,228 reads mapped to the *Ddw1* locus. Two variant calling methodologies were used. The first being a long read specific approach with which 1,918,882 SNPs were identified across the six lines with the statistically significant 5R region (Table 2). 71,140 of these variants were located to the highly significant portion of the locus. Of these only 66 appeared in the putative genic region of *ScGA2ox12*. Given the lack of variants identified within *GA2ox12*, particularly in the dwarf donors which had 0 and 3 variants identified respectively, the alignment in that location was inspected visually. Clear additional variants were visually detected between the dwarf and elite material. Thus, a second variant calling approach, typically used for short reads, was used to supplement this data within *GA2ox12* specifically. The supplementary variant calling identified an additional 125 SNPs (Table 2).

**Table 2.** Summary of ONT sequencing in wildtype elite and dwarf material. The table details the number of variants (SNP only) identified within the significant and highly significant regions of the identified *Ddw1* locus, the number of SNPs that were located within the *GA2ox12* gene model from the primary variant calling methodology and the number of SNPs identified within *GA2ox12* using the supplementary variant calling methodology.

| Genotype | Significant *Ddw1* region | Highly significant *Ddw1* region | SNPs identified in genic *Ddw1* QTL regions | Additional genic SNPs from supplementary approach |
| --- | --- | --- | --- | --- |
| KWL_416 | 506,604 | 18,140 | 60 | 84 |
| KWL_828 | 503,030 | 17,821 | 8 | 45 |
| KWL_1066 | 371,618 | 14,040 | 0 | 19 |
| KWL_1019 | 470,686 | 16,424 | 0 | 16 |
| Donor A (dwarf) | 494,875 | 16,484 | 0 | 57 |
| Donor B (dwarf) | 611,144 | 21,050 | 3 | 51 |
| total | 1,918,882 | 71,140 | 66 | 125 |

Of the 16,541 and 21,101 allelic differences between the reference and donors A and B respectively, 7,314 were found to be unique to Donor A and 12,217 were unique to Donor B (Supplementary Table 1). There were however 1,974 variants found to be shared between donors A and B that differed from the reference genome and the genotypes found in the four representative elite lines (Supplementary Table 1). Furthermore, 297 of the shared dwarf specific SNPs are located within the 24 of the 49 gene models in the highly significant region.

## Discussion

Dwarfism in cereals is a trait of great interest and benefit. However, it remains in its infancy in rye. This study provides steps forward in this topic, offering new variant data that can be easily converted to the KASP platform for tracking *Ddw1*.

Here GWAS analysis was used to confirm the relevant locus as *Ddw1* within the selected dwarf donor material. The observed segregation patterns in two unrelated donors match that of a dominant locus as described by Korzun et al. [7] and Braun et al. [3]. The locus identified, whilst broader than the locus defined by Braun et al. [3], fully encapsulates the Braun et al. *Ddw1* locus and is separate from other identified dominant dwarf loci in rye, located on alternative chromosomes [5, 12, 16]. Thus it can clearly be identified as *Ddw1* rather than a novel or alternative locus.

Previous studies have mapped *Ddw1* to the long arm of chromosome 5R [3]. Here, genetic analysis in the backcrossed populations deriving from donor A identified *Ddw1* as representing the most significant genetic determinant of height, with the most significant markers from the population delimiting a 3.5 Mb interval. This was also the only statistically significant locus identified in this study. The size of the locus identified in this study denotes a region of 49 high confidence genes based on the Lo7 annotation, including some gene models with growth relevant attributes, such as gene models associated with the gibberellic acid cycle and auxin canalisation. Thus, according to the Lo7 annotation, there would be eight genes of potential interest within the *Ddw1* locus in addition to those highlighted in the 2019 study [3]. If this research were to be taken further it would benefit from a supporting genome assembly and subsequent annotation of a dwarf *Ddw1* line. It is possible that the current Lo7 annotation does not include the precise *Ddw1* gene model due to deletion or accumulated mutations to such an extent that the gene is not predicted as a gene model bioinformatically from the genomic sequence, as Lo7 is a tall elite line.

Whilst previous studies have developed markers relating to the *Ddw1* locus, this is the first study to carry out long read genomic sequencing on dwarf rye material. By utilising a selection of representative elite material and dwarf donor material, which were confirmed to as *Ddw1* carriers, 71,140 variants have been identified across the *Ddw1* locus. 21,505 of which were specifically isolated as showing allelic variation association with the dwarf material (Supplementary Table 1), relative to the representative elite lines. Of the variants identified, it was found that 161 are also present amongst the PCR primer intervals of markers developed by Braun et al. [3]. It was found that 18 of the PCR markers from the Braun study reside within the highly significant *Ddw1* locus described here (Table 3). Between the primer pairs of these markers, the number of found variants present in that region were calculated. In contrast to the variants described in this study, the PCR primers [3] do not indicate an allelic state of a SNP. Thus, it cannot be determined from this study alone whether the findings here support or contradict the previous results. Rather these datasets could be utilised in conjunction to further improve the understanding of the *Ddw1* locus.

**Table 3.**
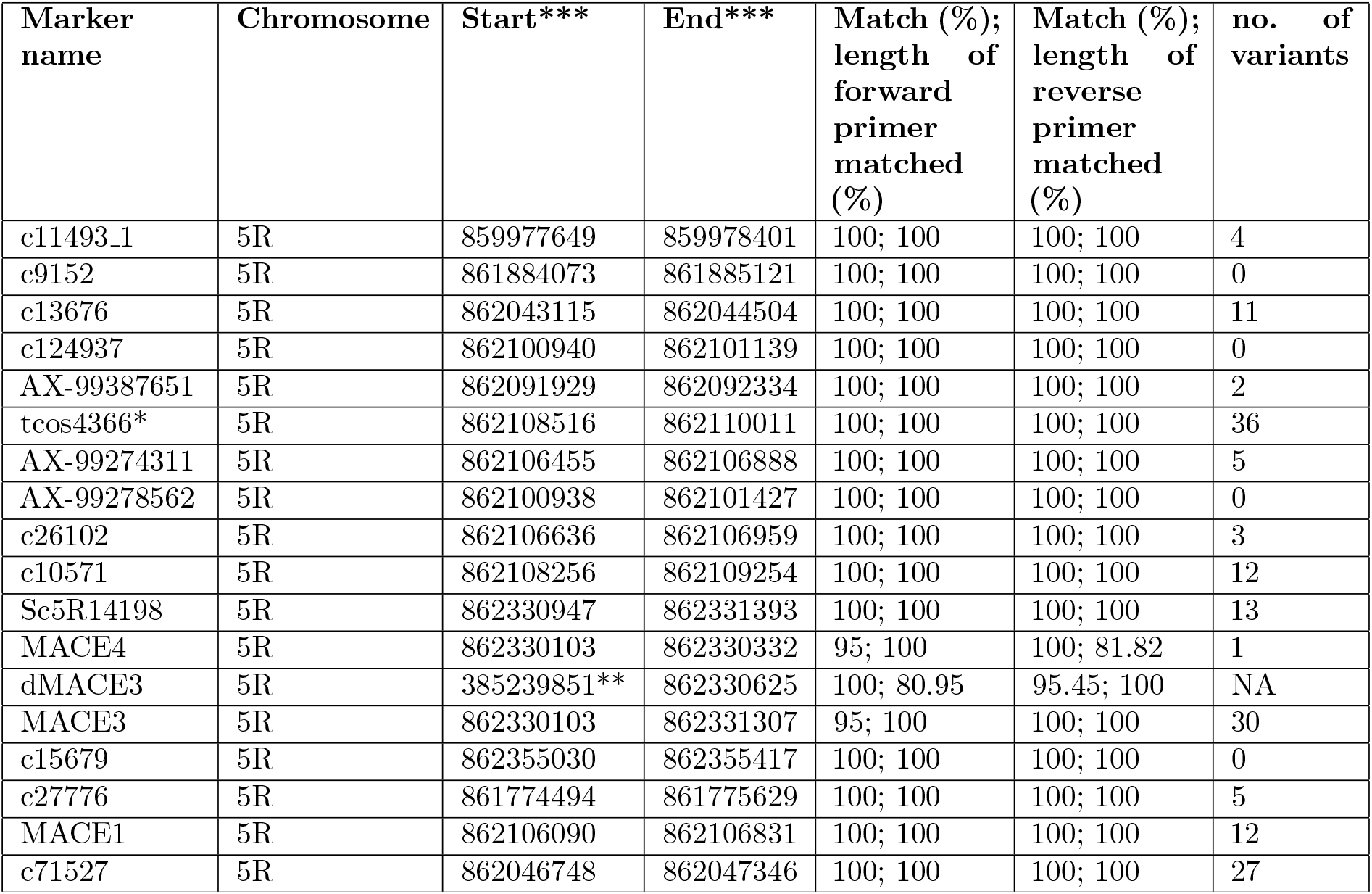
Table of markers developed by Braun et al. (2019) and where they are located physically relative to the Lo7 reference genome. *marker developed by Akhunova et al. [1]. **the forward primer of dMACE3 could not be located within the *Ddw1* locus. ***primer locations (bp) determined with BLAST [18].

A resource as extensive as the variants detailed in this study provides much information for further marker development. For example, this data can be used to find background independent markers, directly associated with dwarfism, or for the development of markers in the flanking regions of the locus to assist in selection against linkage drag. To allow for rapid selection for the allelic state at the *Ddw1* locus, a closely linked KASP genetic marker could be developed from the identified variants. KASP markers represent a useful tool for marker assisted selection for plant height within a breeding program.

## Methodology

Three biparental BC2 populations of rye (S. cereale L.) were developed via two rounds of back-crossing to elite breeding material (non-dwarf). These were phenotyped at EC69. Two of these populations were selected for GWAS. Additional representatives of elite breeding material and two dwarf donors were selected for sequencing (Table 4). The dwarf donors originate from different sources. Donor A is an internal KWL resource acquired from Russia. And Donor B is derived from R347/1 [7]. Genomic DNA of the two Donor A-based BC2 populations was obtained from early leaf material. Genomic DNA of the six lines (Table 2) was obtained via high molecular weight DNA extraction following 72 hours in darkness. Following library preparation, samples of the six lines were sequencing using the PromethION x24 sequencer to coverage of 10X. All samples were sequenced with the SQK-LSK110 library kits on FLO-PRO002 flowcells. The data were basecalled with Guppy v4.2.2 (https://community.nanoporetech.com/) using the “dna r9.4.1 450bps hac prom” model.

**Table 4.** Origin of *Ddw1* donors utilized in this study. Table showing their mean heights (PH) when categorized as dwarf or tall (wildtype).

| Name of Donor | Origin of Donor | Mean PH (cm) dwarf-type | Mean PH (cm) wild-type | Reference |
| --- | --- | --- | --- | --- |
| Donor A | Russia | 60 | 95 | KWL internal |
| Donor B | R347/1 | 83 | 132 | Korzun et al. (1996) |

The selected material was genotyped using an internally designed SNP chip, resulting in 12,908 polymorphic SNPs.

Following genotyping, GWAS analysis was undertaken using a kinship matrix calculated by IBS, implemented with GenAbel [2]. Thresholds of *−log*_10_*P* = 5.3 and *−log*_10_*P* = 15 were termed ‘significant’ and ‘highly significant’, respectively. The above-mentioned genotype data was utilised in conjunction with a physical map of the SNP chip, determined by aligning SNP marker designs to the Lo7 reference genome [14]. Gene models for this region were extracted from the Lo7 annotation [14].

The long read sequences were trimmed to remove adapters, using cutadapt [11], and quality filtered prior to genome alignment to Lo7 using minimap2 [9]. Two variant calling methods were conducted. Firstly, variants on chromosome 5R were called using Longshot [4] with default parameters, which is designed for long read sequences and accounts for the higher error rate found in ONT sequencing. Secondly, the region of the putative candidate gene for *Ddw1* was variant called using bcftools mpileup [10] in order to identify additional SNPs.

## Supplementary Material

**Supplementary Table 1**. Subset of variants at 21,505 genomic positions from highly significant *Ddw1* region that are represented within either of the dwarf donor lines and differ from the reference genome and the other four elite lines.

## References

1. E. D. Akhunov, A. R. Akhunova, O. D. Anderson, J. A. Anderson, N. Blake, M. T. Clegg, and J. Dvorak. Nucleotide diversity maps reveal variation in diversity among wheat genomes and chromosomes. BMC Genomics, 11:1–22, 2010.

2. Y. S. Aulchenko, S. Ripke, A. Isaacs, and C. M. Van Duijn. GenABEL: an r library for genome-wide association analysis. Bioinformatics, 23:1294–1296, 2007.

3. E. M. Braun, N. Tsvetkova, B. Rotter, D. Siekmann, K. Schwefel, N. Krezdorn, and B. Hackauf. Gene expression profiling and fine mapping identifies a gibberellin 2-oxidase gene co-segregating with the dominant dwarfing gene Ddw1 in rye (Secale cereale l.). Frontiers in Plant Science, 857, 2019.

4. P. Edge and V. Bansal. Longshot enables accurate variant calling in diploid genomes from single-molecule long read sequencing. Nature Communications, 10:4660, 2019.

5. Z. Kantarek, P. Masojć, A. Bienias, and P. Milczarski. Identification of a novel, dominant dwarfing gene (Ddw4) and its effect on morphological traits of rye. PLoS One, 13:e0199335, 2018.

6. V. D. Kobylianski. On genetics of the dominant factor of short straw rye. Genetika, 8:12–17, 1972.

7. V. Korzun, A. Börner, and G. Melz. Rflp mapping of the dwarfing (Ddw1) and hairy peduncle (Hp) genes on chromosome 5 of rye (Secale cereale l.). Theoretical and Applied Genetics, 92:1073–1077, 1996.

8. F. Laidig, T. Feike, B. Klocke, J. Macholdt, T. Miedaner, D. Rentel, and H. P. Piepho. Long-term breeding progress of yield, yield-related, and disease resistance traits in five cereal crops of German variety trials. Theoretical and Applied Genetics, 134:3805–3827, 2021.

9. H. Li. Minimap2: pairwise alignment for nucleotide sequences. Bioinformatics, 34:3094–3100, 2018.

10. H. Li, B. Handsaker, A. Wysoker, T. Fennell, J. Ruan, N. Homer, and R. Durbin. The sequence alignment/map format and SAMtools. Bioinformatics, 25:2078–2079, 2009.

11. M. Martin. Cutadapt removes adapter sequences from high-throughput sequencing reads. EMBnet. journal, 17:10–12, 2011.

12. G. Melz. Beiträge zur Genetik des Roggens (Secale cereale l.). Doctoral dissertation, Berlin, 1989.

13. T. Miedaner, M. Hübner, V. Korzun, B. Schmiedchen, E. Bauer, G. Haseneyer, and J. C. Reif. Genetic architecture of complex agronomic traits examined in two testcross populations of rye (Secale cereale l.). BMC Genomics, 13, 2012.

14. M. T. Rabanus-Wallace, B. Hackauf, M. Mascher, T. Lux, T. Wicker, H. Gundlach, and N. Stein. Chromosome-scale genome assembly provides insights into rye biology, evolution and agronomic potential. Nature Genetics, 53:564–573, 2021.

15. R. H. Schlegel. Rye: genetics, breeding, and cultivation. CRC Press, 2013.

16. S. Stojal-owski, B. Myśkśw, and M. Hanek. Phenotypic effect and chromosomal localization of Ddw3, the dominant dwarfing gene in rye (Secale cereale l.). Euphytica, 201:43–52, 2015.

17. T. Würschum, S. M. Langer, C. F. H. Longin, M. R. Tucker, and W. L. Leiser. A modern green revolution gene for reduced height in wheat. The Plant Journal, 92:892–903, 2017.

18. J. Ye, S. McGinnis, and T. L. Madden. BLAST: improvements for better sequence analysis. Nucleic Acids Research, 34:W6–W9, 2006.

